# Adolescent ethanol exposure promotes mechanical allodynia and alters dopamine transmission in the nucleus accumbens shell

**DOI:** 10.1101/2022.09.20.508728

**Authors:** Abigail M. Kelley, Eric J. Del Valle, Samin Zaman, Anushree N. Karkhanis

**Affiliations:** Department of Psychology, Developmental Exposure to Alcohol Research Center, Binghamton University – SUNY, Binghamton, NY, USA

## Abstract

Excessive alcohol consumption in adolescence can disrupt neural development and may augment pain perception. Recent studies have shown that the nucleus accumbens (NAc) shell is involved in mediating pain sensitivity after peripheral inflammation in rodent models of chronic pain and alcohol use disorder (AUD). Interestingly, there have been very few studies examining the impact of chronic ethanol exposure during adolescence on pain sensitivity in adulthood. Therefore, in this project we investigated the impact of adolescent chronic intermittent ethanol (aCIE) exposure on mechanical allodynia and thermal hyperalgesia. Furthermore, given the involvement of the NAc shell in pain processing and chronic ethanol mediated changes, we measured changes in accumbal dopamine kinetics during protracted withdrawal. We found that both male and female aCIE rats show mechanical allodynia during withdrawal; however, only male rats exhibit thermal hyperalgesia during protracted withdrawal. Furthermore, male and female aCIE rats show greater evoked tonic dopamine release, maximal rate of dopamine reuptake, and dopamine affinity to the dopamine transporter in the NAc shell compared to controls. With phasic stimulation, aCIE rats also showed greater dopamine release compared to air exposed rats. These data suggest that aCIE exposure exacerbates pain sensitivity during withdrawal. Furthermore, based on prior literature, it is possible that the increased pain sensitivity may be driven, at least in part, by augmented dopamine kinetics in the NAc shell observed in the current study.

## INTRODUCTION

Over 10% of the adolescent population consuming alcohol either binges or consumes heavy amounts [1]. Binge drinking during this developmentally critical window exacerbates the perception of pain sensitivity during adulthood following abstinence from alcohol [2]. Similarly, rodent studies report that chronic ethanol exposure during adolescence or adulthood augments pain sensitivity to electric tail shock [3], and ethanol exposure during adulthood produces hyperalgesia during withdrawal [4,5]. Furthermore, female rats with prior ethanol history and peripheral inflammation exhibit augmented ethanol consumption [5], suggesting a bidirectional complex relationship between alcohol and pain perception. Both pain and alcohol exposure can also promote neuroadaptations. Preclinical literature highlights overlapping neural circuits that are involved in alcohol use disorder (AUD) and pain, for example, the nucleus accumbens (NAc) is involved in integrating pain and AUD processing [2]. Particularly, acute ethanol exposure in anesthetized adult rats significantly blunts tail pinch-induced terminal dopamine response from the NAc [6], and prior inflammatory pain exposure blunts ethanol-mediated dopamine elevation in the NAc [7]. These data suggest that chronic ethanol exposure could potentially promote hyperalgesia by regulating overlapping neural circuits, specifically with perturbations in the accumbal dopamine system.

While there are a number of rodent studies demonstrating changes in dopamine transmission in the NAc following ethanol exposure in adulthood [8–14], studies examining the effect of adolescent ethanol exposure on accumbal dopamine are relatively few and with disparate results [15–19]. Furthermore, these previous studies have examined dopamine in the NAc core. Surprisingly, to date, only one study has examined dopamine changes in the NAc shell and reported that adolescent ethanol exposure produces a *hyper*dopamine state [19]. It is particularly important to elucidate the impact of adolescent ethanol exposure on dopamine transmission in the NAc shell because in adult rats ethanol-induced elevation in dopamine is greater in the shell compared to core [20]. Moreover, the focus on adolescent ethanol exposure is important given that the dopamine system is maturing during this critical window of development [21,22].

The NAc is also involved in pain processing as accumbal activity was increased at the onset of a painful stimulus in humans [23] and extracellular levels of dopamine in the NAc shell were elevated immediately after foot shock termination in rodents [24]. Similarly, application of capsaicin, a noxious stimulus, attenuated accumbal dopamine tone, but augmented dopamine release evoked by phasic stimulations [25]. Furthermore, the mesolimbic dopamine system is dysregulated as a result of chronic pain and may be highly involved in pain processing resulting in underlying comorbidities highlighted earlier [26].

Despite evidence suggesting greater ethanol consumption during adolescence and later hyperalgesia in humans, very few studies have examined the neural correlates that produce these perceptual changes. In the current study we have established an adolescent chronic intermittent ethanol vapor (aCIE) exposure model and identified mechanical allodynia in males and females, albeit to a greater extent in male rats, while thermal nociception was only apparent in male rats through abstinence. Furthermore, we found that aCIE facilitates dopamine release and uptake kinetics during protracted abstinence in both sexes.

## METHODS

### Subjects

Male (n=32) and female (n=32) Long Evans rats procured from Envigo (Indianapolis, IN) were used in this study. Rats arrived on post-natal day 21 (PD21) and were pair housed with a same-sex cage-mate to acclimate for seven days. Rats were placed on a 12-hour light-dark cycle with lights on at 7:00 and lights off at 19:00. Standard rat chow (LabDiet 5L0D - PicoLab Laboratory Rodent Diet, ScottPharma Solutions, Marlborough, MA) and tap water were provided to the animals ad libitum. **Figure 1A** represents an experimental timeline. All procedures were in compliance with the Institutional Animal Care and Use Committee (IACUC) of the State University of New York at Binghamton.

**FIGURE 1:**
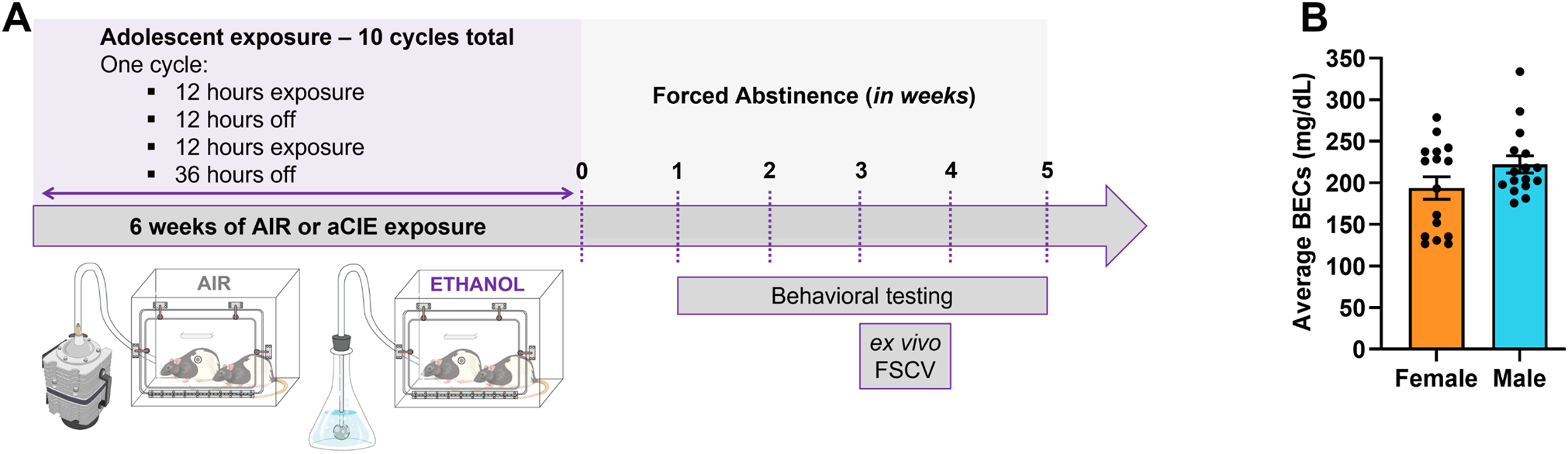
Schematic of experimental timeline and blood ethanol concentrations (BECs). **(A)** Adolescent chronic intermittent ethanol (aCIE) experimental procedure. Male and female rats were exposed to ten cycles of aCIE. Each cycle was two days long with 12 hours of ethanol or air exposure followed by 12 hours of room air after day one and 36 hours of room air after day two, before the next cycle from postnatal day 28 – 65. Following the last exposure, a subset of rats underwent five weeks of behavioral testing. In a different subset of rats, ex vivo fast scan cyclic voltammetry (FSCV) was conducted between weeks three and four of abstinence. **(B)** Blood ethanol concentration (BEC) averaged for each rat from four collections: after cycles one, four, seven, and ten (mean BECs: male rats, 222.2+10.35 mg/dL; female: 193.8+13.51 mg/dL). Vapor chamber settings were adjusted to maintain targeted BECs throughout the experiment.

### Adolescent chronic intermittent ethanol exposure

Rats were exposed to ethanol vapor or air in sealed chambers for two days (12 hours ethanol/air exposure during the dark phase and 12 hours room air) followed by 36 hours abstinence (one cycle) between PD 28 – 65 (**Figure 1A**). Cycles were repeated 10 times. Food was replaced after the first exposure of each cycle, and food and cages were changed after the second exposure within a cycle.

### Blood ethanol concentrations

Tail blood samples were collected immediately after cessation of the first, fourth, seventh, and tenth cycles to measure blood ethanol concentrations (BECs). Samples were then centrifuged and spun at 4°C for 15 minutes at 14,000 RPMs. Serum was separated from plasma and analyzed for ethanol concentration using ANALOX (Model Number AM1; Analox Instruments Ltd. London, UK). BECs were targeted to be 200 mg/dL. Average BECs measured across four cycles (cycles 1, 4, 7, and 10) are shown in **Figure 1B** (Female: 193.8±13.51; Male: 222.2±10.35). The ethanol vapor levels were adjusted to maintain this BEC average throughout the ten cycles.

### Behavioral assessment of mechanical allodynia

Mechanical allodynia was measured using von Frey filaments (58011; Stoelting Illinois, USA) following one, two, three, four, and five weeks of abstinence. We followed a procedure slightly modified from a previous study [27]. Briefly, male and female rats exposed to air or aCIE (n=32; 8 rats per group) were first habituated to the testing apparatus by placing them atop a metal grid (1.27 X 1.27 cm grid size) for 30 minutes each day for two consecutive days. The metal grid allowed complete access to hind paws. On the day of testing, animals were habituated to the apparatus for ten minutes before probing. All behavioral experiments occurred within the first three hours of the light cycle. Rats were probed with calibrated fibers ranging from handle mark values of 1.65 (0.008 g target force) to 6.65 (300 g target force) log stimulus intensity on the plantar area of their right hind paw until they retracted or licked the probed paw. Once the rats responded, they were probed with lower and higher target force fiber to identify the response threshold.

### Behavioral assessment for thermal nociception

Response to thermal nociception was measured after one and four weeks of abstinence in a subset of the rats (n=16; 4 rats per group) tested for mechanical allodynia on separate days using a hot plate (Model Number LE 7406; Panlab, Spain). Thermal nociception assessment was always done after mechanical allodynia assessment. Rats were acclimated to the hot plate apparatus (turned off) for one day for 30 minutes. On the day of testing, rats were habituated to the hot plate apparatus for ten minutes prior to turning on the hot plate to a temperature of 52.4°C + 0.2. The latency to paw withdrawal or licking was recorded. A cutoff of 30 seconds was set to prevent tissue damage to the animal.

### Ex vivo fast scan cyclic voltammetry (FSCV)

We used *ex vivo* FSCV to determine the impact of aCIE on dopamine kinetics in the NAc shell after forced protracted abstinence (3 to 4 weeks) in a separate group of rats that were not subject to behavioral testing. FSCV procedures that we use were similar to those used in previous studies [18,28]. Briefly, rats (n=32; 8 rats per group) were anesthetized with isoflurane and euthanized via rapid decapitation six hours into the dark cycle. Brains were collected promptly and immersed in cold, oxygenated artificial cerebrospinal fluid (aCSF; 126 mM NaCl, 2.5 mM KCl, 1.2 mM NaH2PO4, 2.4 mM CaCl2, 1.2 mM MgCl2, 25 mM NaHCO3, 11 mM glucose, 0.4 mM L-ascorbic acid and the pH was adjusted to 7.4) and then sliced on a vibratome (Leica, VT1200 S, Deerfield, IL, USA). Brain slices (300 μm thick) containing the NAc shell were harvested and transferred to a recording chamber with continuous flow of oxygenated aCSF (32°C). Dopamine efflux was induced using a bipolar stimulating electrode (8IMS3033SPCE, Plastics One, Roanoke, VA, USA) and measured using a carbon fiber recording electrode (~150 μm length, 7 μM radius; C 005722/6, Goodfellow, Huntingdon, England). The recording and stimulating electrodes were placed approximately 100 μm apart. The stimulating electrode was placed on the surface of the slice, while the recording electrode was placed 100 μm below the surface of the slice. Dopamine efflux was induced by a single, rectangular, 4.0-ms duration electrical pulse (750 μA, monophasic, inter-stimulus interval: 300 seconds). To detect dopamine release, a triangular waveform (−0.4 to +1.2 to −0.4V vs. Ag/AgCl, 400V/sec) was applied every 100 ms to the recording electrode. After stable dopamine release was established, multiple pulse stimulations at systematically varying frequencies (2, 5, and 10 pulses at 5, 10, 20, 100 Hz) were used to evoke dopamine release. Recording electrodes used for each experiment were calibrated using 3.0 μM dopamine in order to quantify the concentration of evoked dopamine release. All FSCV recordings were acquired and analyzed using Demon Voltammetry and Analysis software [29]. Dopamine kinetics were determined using the Michaelis-Menten model [30].

### Statistical analysis

All data in this study were analyzed using GraphPad Prism 9 (GraphPad Software, La Jolla, CA, USA). All data are reported as mean + standard error of the mean. Von Frey and hot plate data were analyzed using repeated measures two-way analysis of variance (RM two-way ANOVA). Voltammetry data for baseline dopamine release and rate of uptake were analyzed using RM two-way ANOVA, with sex and exposure set as independent variables. Since no effect of sex was found, male and female data were combined and analyzed using Student’s t-test. Dopamine excitability voltammetry data were first analyzed using three-way ANOVA with sex, exposure, and stimulation parameter as independent variables to assess for sex differences. Since no sex differences were found, data from male and female rats was combined and analyzed using RM twoway ANOVA, with exposure and stimulation parameter as independent variables. RM two-way ANOVA tests were followed by Sidak’s post-hoc pairwise comparisons. The significance level for all statistical measures was set at *p* < 0.05.

## RESULTS

### Chronic intermittent ethanol exposure during adolescence augments touch sensitivity

In order to examine the impact of aCIE exposure on mechanical allodynia, we used von Frey filaments during weeks one through five of abstinence. The experimental timeline is shown in **Figure 2A**. We first conducted an omnibus RM three-way ANOVA with sex, exposure, and abstinence time (repeated measure) as independent variables. This three-way ANOVA revealed a main effect of sex (F _(1,28)_ = 25.91; p < 0.0001), a main effect of exposure (F _(1,28)_ = 28.16; p < 0.0001), and an interaction between exposure and sex (F _(1,28)_ = 4.381; p = 0.0455). Because the sex parameter was significant, we ran separate two-way ANOVAs to first assess sex differences in AIR-exposed male and female rats (**Figure 2B**) and then to assess exposure effects in female (**Figure 2C**) and male (**Figure 2D**) rats separately. A RM two-way ANOVA comparing AIR-exposed male and female rats (**Figure 2B**) revealed a main effect of sex (F _(1,14)_ = 21.60; *p* = 0.0004), while there was no interaction of sex x abstinence time (F _(4,56)_ = 0.7761; *p* = 0.5454) and no main effect of abstinence time (F _(4,56)_ = 0.1391; *p* = 0.9670). Female rats exhibited a lower threshold to touch sensitivity compared to male rats. Next, we compared responsivity on the von Frey test in AIR and aCIE exposed female (**Figure 2C**) and male (**Figure 2D**) rats. A RM two-way ANOVA showed a main effect of exposure in both sexes [female: (F _(1,14)_ = 6.544; *p* = 0.0228), male: (F _(1,14)_ = 22.61; *p* = 0.0003)], but no interaction effect of abstinence time x exposure [female: (F _(4,56)_ = 0.1308; *p* = 0.9705), male: (F _(4,56)_ = 1.836; *p* = 0.1348)] or main effect of abstinence time [female: (F _(4,56)_ = 0.9344; *p* = 0.4508), male: (F _(4,56)_ = 1.920; *p* = 0.1197)]. Overall, rats exposed to aCIE exhibited potentiated touch sensitivity in both sexes, which appeared to persist at least five weeks into abstinence.

**FIGURE 2:**
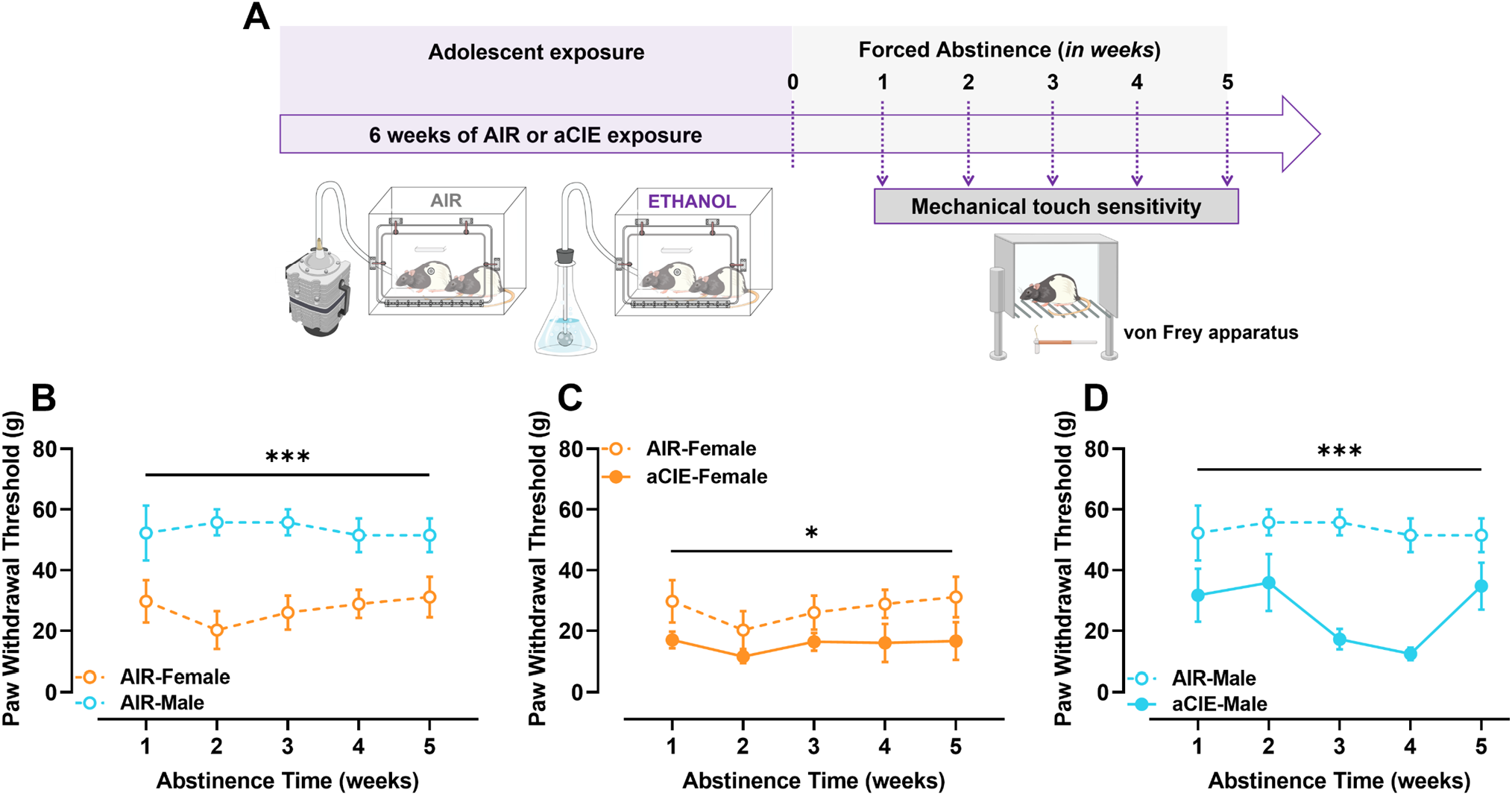
Adolescent ethanol exposure produced mechanical allodynia. **(A)** Mechanical touch sensitivity behavioral testing experimental timeline. Following aCIE exposure, rats underwent five weeks of behavioral testing to measure mechanical allodynia from early to protracted abstinence. **(B)** Air exposed female rats (open orange circles and dotted line) showed greater mechanical touch sensitivity compared to air exposed male rats (open blue circles and dotted line). **(C)** aCIE exposed female rats (closed orange circles and solid line) exhibited greater mechanical touch sensitivity compared to air exposed female rats (open orange circles and dotted line). **(D)** aCIE exposed male rats (closed blue circles and solid line) showed greater mechanical touch sensitivity compared to their air exposed counterparts (open blue circles and dotted line). \**p*< 0.05, \*\*\**p*< 0.001, main effect of sex (B) and exposure (C and D).

### Exposure to aCIE produces thermal nociception in male rats only

In order to determine the effects of aCIE exposure on thermal nociception, we used a hot plate set to 52.4°C during weeks one and four of abstinence (**Figure 3A**). Similar to the von Frey assay, to assess sex differences in thermal nociception, we first conducted an omnibus RM three-way ANOVA, which revealed a significant interaction between sex and abstinence time (F _(1,12)_ = 5.025; *p* = 0.0447) and sex, exposure, and abstinence time (F _(1,12)_ = 5.223; *p* = 0.0413). Because we found these interactions, we performed separate RM two-way ANOVAs to assess sex differences in AIR-exposed male and female rats (**Figure 3B**), and then exposure effects in female (**Figure 3C**) and male (**Figure 3D**) rats separately. A comparison of paw withdrawal latency in AIR exposed male and female rats (**Figure 3B**) using a RM two-way ANOVA revealed an interaction between abstinence time and sex (F _(1,6)_ = 21.68; *p* = 0.0035), while there was no main effect of abstinence time (F _(1,6)_ = 2.212; *p* = 0.1875) or sex (F _(1,6)_ = 2.227; *p* = 0.1862). Pairwise comparisons using Sidak’s post-hoc analysis showed that female rats exhibited a significantly lower withdrawal latency at the four-week timepoint (*p* = 0.0125). Next, we compared paw withdrawal latency in AIR and aCIE exposed female rats (**Figure 3C**). A RM two-way ANOVA showed no main effects of abstinence time (F _(1,6)_ = 3.708; *p* = 0.1025) or exposure (F _(l·6)_ = 1.823; *p* = 0.2257), or an interaction between abstinence time and exposure (F _(1,6)_ = 0.008914; *p* = 0.9279). Lastly, we compared paw withdrawal latency in AIR and aCIE exposed male rats (**Figure 3D**). A RM two-way ANOVA showed an interaction between abstinence time and exposure (F _(1,6)_ = 11.71; *p* = 0.0141), but no main effects of abstinence time (F _(1,6)_ = 1.429; *p* = 0.2770) or exposure (F _(1,6)_ = 2.730; *p* = 0.1496). Pairwise comparisons using Sidak’s post-hoc analysis showed that aCIE-exposed male rats exhibited a significantly lower paw withdrawal latency at the four-week abstinence timepoint when compared to AIR-exposed male rats (*p* = 0.0130). Post-hoc comparison between week one and four of abstinence revealed a significant difference in AIR-exposed male rats (*p* = 0.0340). These data suggest that aCIE produced thermal nociception selectively in male rats, but only during protracted withdrawal. Furthermore, both aCIE-exposed male rats and AIR- and aCIE-exposed female rats reduced response latency when tested for a second time (4^th^ week of abstinence).

**FIGURE 3:**
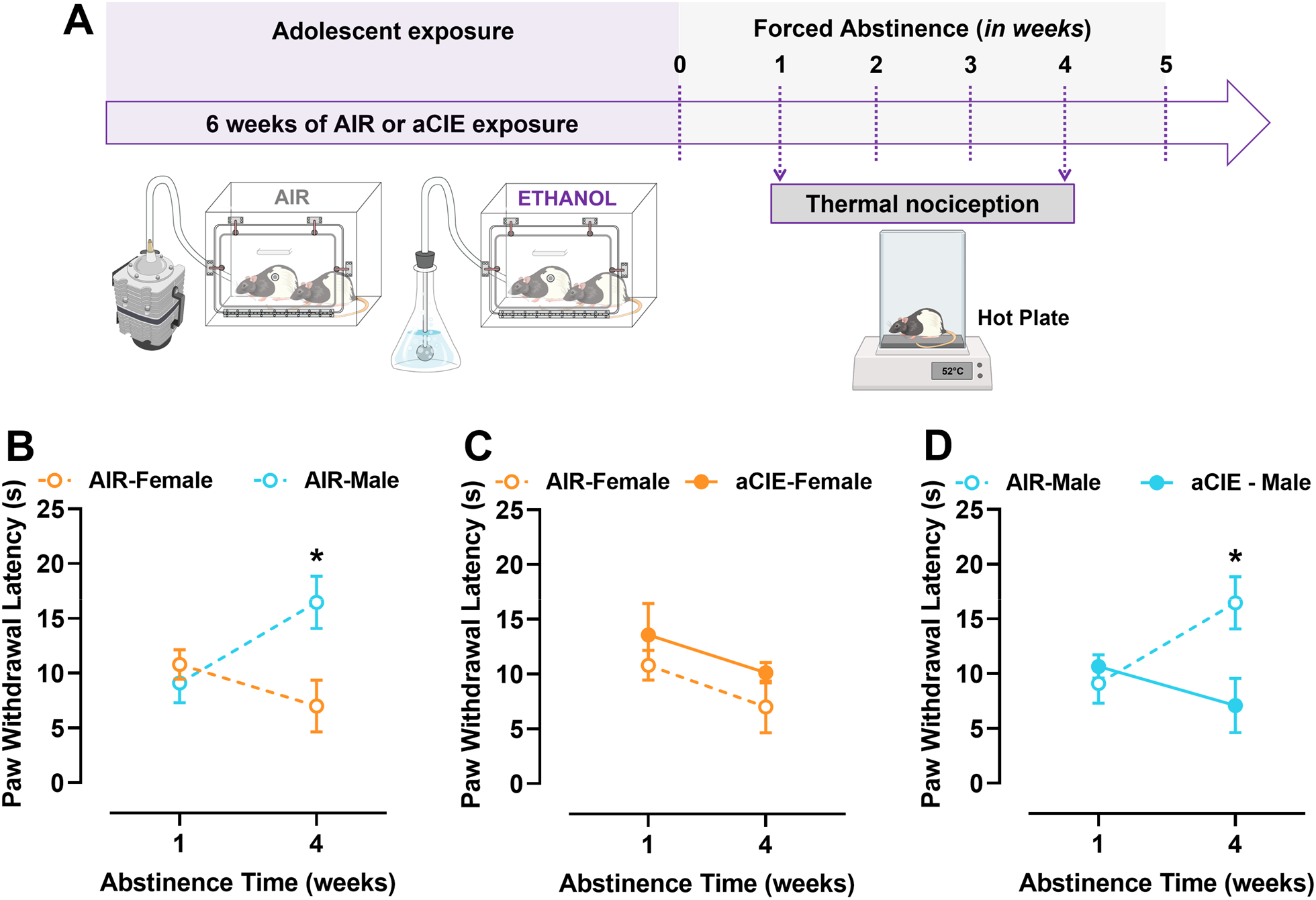
Adolescent ethanol-associated sensitivity to thermal stimuli is observed selectively in male rats. **(A)** Thermal nociception behavioral testing experimental timeline. Following aCIE exposure, rats underwent behavioral testing during weeks one and four of abstinence to measure thermal nociception during early and protracted abstinence using the Hot Plate apparatus. **(B)** Air exposed female rats (open orange circles and dotted line) responded at a lower paw withdrawal latency during week four of abstinence compared to air exposed male rats (open blue circles and dotted line). No difference in responsivity between air exposed male and female rats was observed during week one of abstinence. **(C)** Air (open orange circles and dotted line) and aCIE (closed orange circles and dotted line) exposed female rats exhibited similar thermal nociception during week one and four of testing. **(D)** While no differences were observed between air (open blue circle and dotted line) and aCIE (closed blue circle and dotted line) exposed male rats during week one of abstinence, aCIE males respond at a lower paw withdrawal latency during week four of abstinence. There is no difference during week one of abstinence. \**p*< 0.05, Sidak’s pairwise comparison between sex (B) and exposure (D). aCIE, adolescent chronic intermittent ethanol.

### Chronic adolescent intermittent ethanol exposure perturbs dopamine kinetics

Chronic ethanol exposure in adult rats alters dopamine transmission in the NAc and recent studies have elucidated that NAc shell is involved in regulating pain perception. Because the effect of mechanical sensitivity was consistent through several weeks of protracted abstinence, we measured aCIE mediated changes in dopamine transmission in the NAc shell at weeks 3-4 of abstinence using *ex vivo* FSCV (**Figure 4A**). Recording locations in slices containing the NAc shell are shown in **Figure 4B**; we targeted the medial dashed – AIR, solid – aCIE) rats are shown in **Figure 4C**.

**FIGURE 4:**
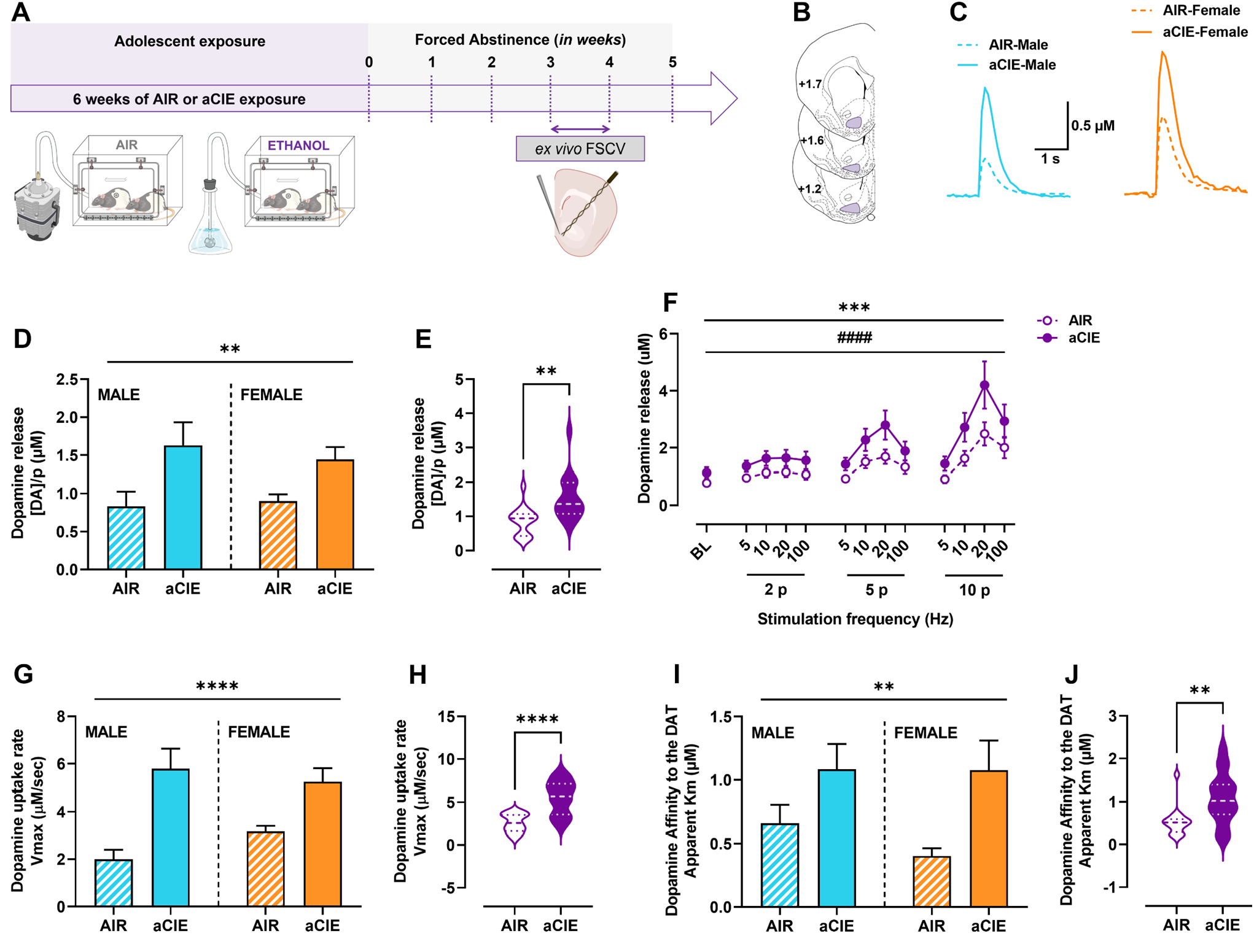
Adolescent ethanol exposure augments dopamine transmission during protracted abstinence. **(A)** *Ex vivo* FSCV experimental timeline; FSCV experiments were conducted between weeks three and four of abstinence following cessation of air or aCIE exposure. **(B)** Coronal sections showing area from which dopamine kinetics were measured (purple shaded region - middle NAc shell) in air and aCIE exposed male and female rats. **(C)** Representative transient dopamine response to single pulse electrical stimulation in male (air, blue dotted line; aCIE, blue solid line) and female (air, orange dotted line; aCIE, orange solid line) exposed rats. These traces show that evoked dopamine response was significantly greater in male and female aCIE exposed rats compared to their air exposed counterparts. **(D)** Tonic dopamine release was significantly greater in NAc shell of aCIE exposed male (blue solid bar) and female (orange solid bar) compared to air exposed rats (male, blue shaded bar; female, orange shaded bar). **(E)** Comparison between and air (open violin plot) and aCIE (closed violin plot) exposed rats combined across sexes showed greater dopamine release in NAc shell of aCIE exposed rats. The violin plots show the distribution of data points; vertical aspect of the plot shows variability, and the horizontal aspect shows population size at a given response level. Dashed lines in the violin plot represents means and the dotted line above and below the dashed line represent quartiles. **(F)** Phasic dopamine release in the NAc shell of aCIE exposed rats (male and female data combined, solid circles and lines) was significantly greater than their air exposed counterparts (male and female data combined, open circles and lines). **(G)** Dopamine uptake rate was significantly greater in aCIE compared to air exposed male (air, shaded blue bars; aCIE solid blue bars) and female (air, shaded orange bars; aCIE, solid orange bars) rats. **(H)** aCIE (solid violin plot) exposure promotes faster dopamine uptake in male and female rats compared to air (open violin plot) exposure. All characteristics of the violin plot are same as in panel E. **(I)** Dopamine affinity to the DAT was significantly greater in male and female rats exposed to aCIE compared to those exposed to air. Bar characteristics same as in panels D and G. **(J)** aCIE exposure augments dopamine affinity to the DAT. All violin plot characteristics are same as those in panel E. \*\**p*< 0.01, \*\*\**p*< 0.001, \*\*\*\**p* < 0.0001, main effect of exposure. ^####^*p*<0.000**1**, main effect if stimulation parameter. FSCV, fast scan cyclic voltammetry; aCIE, adolescent chronic intermittent ethanol exposure; DAT, dopamine transporter.

#### Chronic adolescent intermittent ethanol exposure potentiates dopamine release

To investigate the effects of aCIE exposure on dopamine release, we measured evoked dopamine responses following electrical stimulations applied at single pulse (tonic, **Figure 4D and E**) and multiple pulses with varying frequencies (phasic; **Figure 4F**). We quantified tonic dopamine release using the Michaelis-Menton equation, which corrects for continuous dopamine uptake that occurs during synaptic transmission. A RM two-way ANOVA with sex and exposure as dependent variables (**Figure 4D**) revealed a main effect of exposure (F _(1,28)_ = 11.03; *p* = 0.0025), but no effect of sex (F _(1,28)_ = 0.0813; *p* = 0.7776) or an exposure X sex interaction (F _(1,28)_ = 0.3900; *p* = 0.5374). Since there was no effect of sex, we subsequently combined male and female data and conducted a Student’s *t*-test comparing dopamine release in AIR and aCIE exposed rats (**Figure 4E**). Tonic dopamine release was significantly greater in aCIE exposed rats compared to AIR exposed counterparts (mean: AIR, 0.866±0.104; aCIE, 1.537±0.167; *t_30_* = 3.409; *p* = 0.0019).

To assess the effect of aCIE on / dopamine release in male and female rats, we conducted a three-way ANOVA with sex, exposure, and stimulation parameter as independent variables and dopamine release as the dependent variable. This analysis revealed a main effect of stimulation parameter (F _(1.361, 38.10)_ = 36.87; *p* < 0.0001) and an interaction between exposure and stimulation parameter (F _(12, 336)_ = 2.987; *p* = 0.0006). With no significant effect of sex, we combined the male and female to analyze it further. Assessment of phasic dopamine release using a RM two-way ANOVA (**Figure 4F**; data from male and female rats combined; exposure and stimulation parameter as dependent variables) revealed an interaction between exposure and stimulation parameter (F _(12,360)_ = 2.904; *p* = 0.0007) and a significant main effect of stimulation parameter (F _(1,30)_ = 2.964; *p* = 0.0955), but no main effect of exposure (F _(1,28)_ = 0.0813; *p* = 0.7776). In summary, aCIE exposure potentiated tonic and phasic dopamine release in a sex independent manner.

#### Exposure to aCIE facilitates dopamine uptake kinetics

Next, we quantified maximal rate of uptake (Vmax) of dopamine by the dopamine transporters (DATs) using Michaelis-Menten equation. A two-way ANOVA assessing sex and exposure as independent variables (**Figure 4G**) revealed a main effect of exposure (F _(1,28)_ = 27.79; *p* < 0.0001), but no effect of sex (F _(i,28)_ = 0.3170; *p* = 0.5779) or an interaction between the two (F _(l·28)_ = 2.415; *p* = 0.1314). Since there was no significant effect of sex, we combined the data from male and female rats within each exposure group and conducted a Student’s *t*-test (**Figure 4H**). Rats exposed to aCIE exhibited significantly greater maximal rate of dopamine reuptake compared to AIR exposed controls (mean: AIR, 2.586±0.271; aCIE, 5.521±0.494; *t*_30_ = 5.208; *p* < 0.0001).

Given the robust effect of Vmax, we quantified Km, which is a measure of the affinity of dopamine to the DAT. A two-way ANOVA assessing sex and exposure effects (**Figure 4I**) revealed a main effect of exposure (F _(1,28)_ = 10.09; *p* = 0.0036), but no effect of sex (F _(1,28)_ = 0.5958; *p* = 0.4467) or an interaction between the two factors (F _(1,28)_ = 0.5914; *p* = 0.4771). Because there was no effect of sex, we combined the data across the two sexes and conducted a Student’s *t-*test assessing the effect of exposure on Km (**Figure 4J**). As with Vmax, aCIE exposed rats exhibited a significantly greater Km compared to their AIR exposed counterparts (mean: AIR, 0.532±0.084; aCIE, 1.082±0.148; *t*30 = 5.208; *p* < 0.0001).

## DISCUSSION

In the current study, we showed that aCIE exposure augmented pain sensitivity and altered dopamine dynamics in the NAc shell. Both male and female aCIE exposed rats exhibited mechanical allodynia. However, only male rats expressed thermal nociception in the hot plate test. In addition, aCIE exposure potentiated tonic and phasic dopamine release, and facilitated maximal rate of dopamine uptake and the affinity of dopamine to the DAT.

Our data showing greater pain responsivity (mechanical allodynia and thermal nociception) in AIR-exposed female compared to male rats are congruent with previous findings in both human and rodent studies. For example, one study found that women have a higher prevalence of chronic pain states and a greater pain sensitivity compared to men [31,32]. Similarly, a rodent study showed that female mice exhibit greater mechanical allodynia and cold hypersensitivity compared to male mice in response to an intrathecal injection of Gp120ADA, an HIV-viral protein [33]. Interestingly, this study further showed that ovariectomized female mice exhibited lower pain sensitivity and cold hypersensitivity compared to female mice with intact ovaries. These data suggest that gonadal hormones likely play a role in shaping sensitivity to pain. Indeed, estradiol has been shown to regulate pain sensitivity as male rats injected with estradiol exhibited greater paw licking after formalin injection into the paw compared to saline controls [34]. In contrast, evidence suggests that testosterone plays a protective role in adjuvant-induced arthritis in male rats [35]. Furthermore, when supraphysiological levels of testosterone were administered to rats of both sexes, paw licking duration in response to formalin decreased for only females [36]. These data suggest that a threshold level of testosterone, for example, basal levels in males, is required for lower pain sensitivity response. Thus, the sex differences that occur in pain sensitivity may be ultimately driven by differences in gonadal hormone levels.

In addition to the inherent sex differences, our study showed that both male and female aCIE-exposed rats responded with paw withdrawal at a lower target force in the von Frey mechanical allodynia test. Our data concur with a recent study showing moderate ethanol exposure during early adolescence (PD 25 – 53) mediated reduction in paw withdrawal threshold during protracted withdrawal in male mice [37]. Furthermore, in the current study, measurements of sensitivity to thermal nociception during protracted withdrawal revealed thermal hyperalgesia selectively in aCIE exposed male rats. In accordance with our data, a recent study reported attenuation in paw withdrawal latency in response to noxious thermal stimulus in male mice during protracted abstinence following adolescent ethanol exposure (PD 25 – 53) [37]. Previous studies have also reported heightened thermal hyperalgesia during withdrawal from chronic ethanol exposure in adulthood in male rats [4,38]. To date, there have been no studies examining thermal nociception in female rodents following ethanol exposure at any age. In the current study, while the AIR exposed male rats exhibited greater response latency when tested in week four for the second time (first time being in week one), aCIE exposed male rats did not show a difference between weeks one and four. This could potentially imply that aCIE exposure blunts neural plasticity in response to thermal nociception stimuli. This is an interesting direction of research for future studies.

The relationship between alcohol and pain may be driven by changes in the NAc shell, as this brain region lies at the intersection of the pain processing and affect- and addiction-related circuits [2,28,39]. A recent study showed that chronic adolescent intermittent ethanol exposure (PD 28 – 48) augments subsequent ethanol-mediated increase in extracellular levels of dopamine in the NAc shell during protracted abstinence in adulthood, suggesting that adolescent ethanol exposure promotes hypersensitivity of the dopamine system [19]. Another study showed greater basal dopamine levels measured using quantitative microdialysis in rats given systemic injections of ethanol during adolescence (PD 35 – 50) followed by 15 days of abstinence compared to saline controls [15], again suggesting adolescent ethanol and withdrawal are associated with enhancement of dopamine transmission. Our study is the first showing greater stimulated tonic and phasic dopamine release in the NAc shell of aCIE exposed rats (male and female) compared to AIR exposed controls during protracted abstinence. The greater stimulated release observed in the current study potentially explains the higher levels of ethanol-induced dopamine quantified in the previous studies. It is important to note that total dopamine tissue content was observed to be lower after five days of abstinence following ethanol exposure between PD 30 and 45 compared to controls [17]. These opposing results could be explained by the difference in withdrawal timepoint - acute abstinence as opposed to the protracted abstinence period used in the current study [15]. Collectively these data suggest that adolescent ethanol exposure may affect dopamine system maturation resulting in altered dopamine transmission in adulthood, but this ultimate change in dopamine transmission is possibly a function of the length of abstinence period.

In addition to changes in release, our current data show an aCIE-associated potentiation of rate of dopamine clearance from the synapse in both male and female rats. This augmented rate of uptake may be driven by enhanced DAT functionality or greater density of DATs. Indeed, one study showed a repeated ethanol exposure associated increase in DAT binding sites in mice [40], and faster rate of uptake was previously shown to be associated with greater DAT expression levels following adolescent psychosocial stress exposure [41]. It is also possible that aCIE exposure increases binding cites at the DATs. This could potentially affect the binding affinity of dopamine to the DAT, which is measured by Km. Interestingly, our current data show potentiated Km in aCIE compared to AIR exposed in both sexes. Thus, in general, we report an aCIE exposure-associated enhancement in dopamine transmission in the NAc shell during protracted abstinence in both male and female rats. Dopamine maturation and pruning occur later in adolescence; i.e., dopamine cell firing peaks at approximately PD 45 (mid-adolescence) followed by pruning of excess synapses during the latter half of adolescence (PD 45 - 65) resulting in reduced firing rates [21,22]. It is possible that aCIE exposure facilitates greater dopamine neuron firing as ethanol is known to increase cell firing of tegmental dopamine neurons [42]. The continuous exposure to ethanol may disrupt the pruning process later in adolescence, arresting the system in a mid-adolescent state.

Withdrawal from ethanol is stressful [43]. Because our aCIE exposure protocol includes forced abstinence periods, it is possible that the rats experience withdrawal-induced stress. Previous studies have shown that chronic social isolation stress during adolescence (age matched to the current study) potentiates dopamine release and clearance rates in the NAc shell and core [28,41,44]. These studies have hypothesized that the potentiated dopamine release is driven by enhanced excitability of dopamine neuron terminals measured using *ex vivo* FSCV [44]. Interestingly, external stimuli such as systemic ethanol administration or nociceptive stimuli such as footshock elevate extracellular levels of dopamine more in socially isolated compared to group housed rats [45]. Thus, it is possible that the potentiated dopamine transmission observed in the current study may contribute to the greater perception of mechanical touch in both sexes.

In summary, the results of this study demonstrated that both male and female rats exposed to chronic intermittent ethanol exposure during adolescence experienced mechanical allodynia during abstinence. However, only males exhibited thermal nociception during protracted withdrawal. Furthermore, we observed greater dopamine reuptake in both male and female aCIE rats compared to their respective controls. In addition, dopamine terminal excitability was greater in aCIE exposed rats. Together, these data suggest that aCIE exposure augments pain sensitivity during abstinence and that this increased pain sensitivity may be driven, at least in part, by augmented dopamine transmission in the NAc shell. The results of the current study have high clinical relevance. Opioid misuse in our society is currently on an uprise, but interestingly it is eminent in a subset of our population. Individuals with a history of alcohol misuse are more likely to misuse opioids [46–48]. For most, the initial use of opioids begins in the clinic to treat pain. Enhanced pain perception and sensitivity, and chronic pain disorders are largely mediated by prior history of ethanol consumption [46]. Thus, individualized medication targeting the dopamine system may help in reducing the opioid dose needed to treat pain particularly in individuals with a history of alcohol misuse during adolescence. In future studies, we will investigate the direct relationship between dopamine transmission and pain regulation to determine mechanisms that may be involved in pain perception during abstinence following aCIE exposure.

## ACKNOWLEDGEMENTS

This study was supported by grants from the NIAAA: P50 AA017823 (Developmental Exposure Alcohol Research Center) and R01 AA028228 (ANK). None of the authors have any conflict of interest to report.

